# Predicting Novel Metabolic Pathways through Subgraph Mining

**DOI:** 10.1101/123877

**Authors:** Aravind Sankar, Sayan Ranu, Karthik Raman

## Abstract

The ability to predict pathways for biosynthesis of metabolites is very important in metabolic engineering. It is possible to mine the repertoire of biochemical transformations from reaction databases, and apply the knowledge to predict reactions to synthesize new molecules. However, this usually involves a careful understanding of the mechanism and the knowledge of the exact bonds being created and broken. There is clearly a need for a method to rapidly predict reactions for synthesizing new molecules, which relies only on the structures of the molecules, without demanding additional information such as thermodynamics or hand-curated information such as atom-atom mapping, which are often hard to obtain accurately.

We here describe a robust method based on subgraph mining, to predict a series of biochemical transformations, which can convert between two (even previously unseen) molecules. We first describe a reliable method based on subgraph edit distance to map reactants and products, using only their chemical structures. Having mapped reactants and products, we identify the reaction centre and its neighbourhood, the reaction signature, and store this in a reaction rule network. This novel representation enables us to rapidly predict pathways, even between previously unseen molecules. We also propose a heuristic that predominantly recovers natural biosynthetic pathways from amongst hundreds of possible alternatives, through a directed search of the reaction rule network, enabling us to provide a reliable ranking of the different pathways. Our approach scales well, even to databases with > 100,000 reactions. A Java-based implementation of our algorithms is available at https://github.com/RamanLab/ReactionMiner

**CCS CONCEPTS:** •Information systems →Data mining; •Applied computing →Bioinformatics;

## 1 INTRODUCTION

Metabolic networks have been curated for hundreds of organisms, with varying degrees of detail and confidence, in popular databases such as the Kyoto Encyclopedia of Genes and Genomes (KEGG; [15]), MetaCyc [5] and MetaNetX [11]. These curated biochemical reaction databases represent the repertoire of biochemical conversions that known enzymes can catalyse. Enzymes, while being remarkably specific, also demonstrate the ability to convert a family of related substrates (e.g. alcohols), to a family of related products (e.g. aldehydes). An important challenge in metabolic engineering is the biosynthesis of novel molecules through heterologous expression of enzymes from other organisms. The ability to perform this *retrosynthesis* of novel molecules hinges on our ability to understand and generalise the abilities of the enzymes, in terms of the chemical reactions that they can catalyse and the substrates that they can act on.

Further, a deeper understanding of the biochemical transformations happening in metabolic networks can shed light on various fundamental questions in biology. For example, are there alternate ways to synthesize common central metabolites such as pyruvate? Why do cells prefer a particular pathway for the conversion of a metabolite such as glucose, to say, pyruvate (glycolysis)? There are also many knowledge gaps in our understanding of microbial metabolism; for example, there are a number of compounds known to be present in microbes, but the exact sequence of reactions and intermediates involved in their biosynthesis remain unknown. It is possible to bridge these knowledge gaps through a careful analysis of the metabolic networks, as we describe herein.

Since the seminal work of Corey and Wipke [9], a number of algorithms have been developed to analyse (bio)chemical reaction networks, to predict pathways and novel routes for metabolite synthesis [3, 4, 6, 13, 17, 20, 24, 29, 30]. For reviews, see [12, 23]. Despite the availability of a wide array of reaction prediction methods, all of them rely on the existence of query molecules in the reaction knowledge-base (“known” molecules in training data). ReactionPredictor [6, 17] is one exception as it can predict reactions for unknown molecules, but it is limited to specific classes of reactions due to its reliance on hand-curated rules. In this work, we present for the first time to the best of our knowledge, a general and fully-automated method for predicting reactions between unknown (previously unseen) molecules. We do so by automatically learning biochemical transformation rules involving substructures of molecules from the reaction knowledge-base and searching for matching substructures in the unseen query molecule, both via subgraph mining techniques. The result is a scalable method that can be efficiently applied to predict novel metabolic routes in thousands of organisms.

Notably, compared to previous methods, which use KEGG atom types [26] or atom–atom mapping information [20, 29], we use no more information than the metabolic reaction database and the chemical structures of the participating molecules. We also demonstrate two important applications of our method: first, we show how our method can be used to identify/recover biochemically preferred pathways between metabolites. Second, we show how pathways to known and novel/unseen compounds can be rapidly predicted. Our approach is very efficient, completely automated, scalable and performs with a higher degree of accuracy compared to state-of-the-art methods.

## 2 RELATED WORK

We now discuss how the previous approaches meet only a subset of the challenges mentioned above. The proposed technique is the first to overcome all of the above challenges. The earliest work [21] focuses on using stoichiometric constraints to identify feasible pathways, where reactions are classified as either being allowed, required or excluded from the pathways. Rahnuma [24] employs a hypergraph model to represent a network between molecules for the prediction and analysis of pathways. An edge connecting two molecules denote that it is possible to convert one to the other. Metabolic Tinker [22] is an open source web-server that uses the entire Rhea database to rank possible paths, based on thermodynamics. All the above techniques, however, fail to generalise for unknown query molecules. PathPred [26] uses a limited number of (Reactant, Product) pairs to predict pathways for a small subset of molecules. However, these pairs and their structural transformations are hand-curated and consequently, the technique is limited to a small collection of reactions. In our technique, we automatically learn both the pairing and the structural transformations.

EC-BLAST [29] proposes an algorithm to automatically search and compare enzyme reactions. Though their approach characterises reactions using patterns derived from atom–atom mappings, they use additional chemical knowledge such as bond energies and do not address our precise problem of predicting chemical reactions. Furthermore, information on bond energies is not readily available. Kotera and co-workers [18] developed a method to learn enzymatic reaction likeness from metabolic reaction databases using chemical fingerprints. From Metabolite to Metabolite (FMM; [7]) is a tool for predicting pathways based on the KEGG. RouteSearch [20] is a recent method to predict pathways using the MetaCyc database. This technique uses atom–atom mappings to search a metabolic network obtained from MetaCyc [5]. Another very recent tool is Metabolic Route Explorer (MRE; [19]), which can rapidly predict pathways in several organisms and rank the pathways via a nice web interface. However, none of FMM, RouteSearch or MRE can predict on unseen molecules.

## 3 METHODS

Fig 1 presents the pipeline of our reaction prediction algorithm. We represent each molecule as a graph, where atoms correspond to vertices and bonds correspond to edges. Given a database of metabolic reactions, we use an effective mapping method based on subgraph edit distance [14] to accurately map transformed metabolites in a reaction. Through graph mining, we then identify the specific subgraph within a graph (molecule) that is critical for a reaction to occur. We call these subgraphs the *reaction signatures*. For example, consider an alcohol to aldehyde conversion (see Fig 2a), where *RCH_2_OH* is converted to *RCHO*, by the enzyme alcohol dehydrogenase. We consider the subgraph corresponding to *CH_2_OH* as the reaction signature, since the rest of the molecule remains un-affected. We then analyze the reaction signatures and characterize the changes they undergo during a reaction and summarize them as *reaction rules*. Connecting back to our example, the reaction rule in this case is *CH_2_OH* changing to *CHO*. All reaction rules that are learned from the database are next consolidated in the form of a *reaction rule network* (RRN). In the RRN, each node is a reaction rule and two rules are connected by an edge if they can potentially form a reaction pathway. This completes the offline phase. In the online phase, given a query to find a pathway from molecule *A* to *B*, we analyze the structures of both *A* and *B* based on the reaction signatures they contain. From this analysis, *A* is mapped to a set of source nodes, and *B* is mapped to a set of destination nodes, in the RRN. Consequently, the prediction problem reduces to finding (optimal) paths between the source and destination nodes in the RRN.

**Figure 1:**
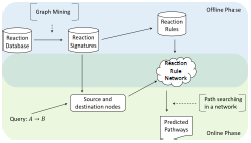
Pipeline of the reaction prediction algorithm. The figure outlines both the offline and online phases of the algorithm. The offline phase involves graph mining of the reaction database to identify reaction signatures, from which reaction rules are subsequently identified and embedded in a reaction rule network (RRN). In the online phase, we search the RRN and predict suitable pathways, on the arrival of a query *A → B*.

### 3.1 Problem Formulation

In this section, we formulate our prediction problem and define the concepts and notations central to our work. We represent each molecule as an undirected graph. A graph *g*(*V, E*) is composed of a set of vertices *V* = {*υ_1_,…,υ_n_}* and a set of edges *E* = {*e* = (*υ_i_, υ_j_*) | *υ_i_, υ_j_* ∈ *V*}. Each vertex and edge have labels denoted *l*(*υ*) and *l*(*e*) respectively. The *size* of a graph is |*E*|. Fig 2b shows the graph representation of a molecule. Atoms correspond to vertices, bonds correspond to edges and bond orders correspond to edge labels.

**Figure 2:**
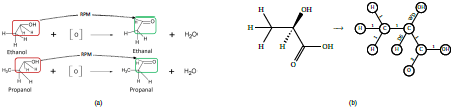
A simple illustration to motivate our approach. (a) Conversion of ethanol and propanol (alcohols) to ethanal and propanal (aldehydes) respectively. Vertices without explicit labels represent Carbon atoms. Notice that although the reactions involve different molecules, the changes (highlighted in red and green boxes) are identical. (b) Representing D-Lactic acid as a graph. Note that double bonds are indicated by a changed edge label, as are *wedges* and *dashes* that represent bond stereochemistry.

The input to our problem is a dataset of chemical *reactions* ℝ. A reaction 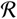 contains two sets of graphs (or molecules): the first set contains the *reactants* and the second set contains the *products* synthesized. We use *RS*(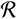) to denote the reactant set in 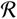 and *PS*(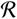) to denote the products. A *pathway P(A, B*) from a molecule *A* to *B* is a chain of reactions 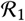,…,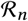 such that *A* ∈ *RS*(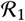), *B* ∈ *PS*(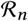), and there is at least one metabolite shared between the product set of one reaction and the reactant set of the next. An example of a pathway is from ethanol to ethanoic acid. Ethanol can be oxidised to form ethanal, and then ethanal can be oxidised to form ethanoic acid.

We now define the pathway prediction problem as follows: Given a training database of reactions (and the structures of the constituent molecules), learn a prediction model 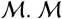 should support the prediction query *Q*(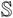, *T*), where 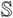 is a (set of) source molecule(s) and *T* is the target molecule. Given this query, 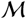 should produce a pathway *P*(*A, T*) where *A* ∈ 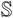. An important aspect of our formulation is that we do not make any assumption of the source or the target molecules being present in the reaction database. The only information we use to learn the prediction model are the structures of the molecules, which is easily available.

### 3.2 Mining Reaction Patterns

Our goal in this section is two-fold. First, we identify the *reaction patterns* existing in the training database. Second, for any given molecule in the reactant set, we should be able to predict the patterns that are applicable on the reactant. To understand what a pattern is in our context, let us revisit Fig 2a. We claim that both reactions follow the same pattern because: (i) in both the alcohol molecules, the exact same subgraph (highlighted in red) is affected, while the remaining portions remain unaltered, (ii) the affected subgraphs undergo an identical change and (iii) the oxidising agent undergoes an identical change to form a water molecule.

In other words, if the same structural change happens in one or more reactions, then that is a pattern. To quantify the *structural change*, we first need to construct a mapping between the graphs in the reactant set to those in the product set. More specifically, the alcohol molecules should be mapped to the aldehyde molecules and the oxidizing agent should be mapped to water. The comparison in the structure of the mapped molecules allows us to quantify the change. We call this operation *reactant–product mapping (RPM)* and use the notation *RPM* (*A, B*) to denote that a reactant *A* has been mapped to a product *B* of the reaction. Clearly, a wrong mapping (such as mapping alcohol to water) would produce spurious results. As we demonstrate later, our RPM allows us to reliably compute meaningful pathways. This is similar to the RPAIR concept used in KEGG and by MRE, but we compute it only using the molecule structures, without resorting to the use of atom–atom mapping information, or even atom types.

Clearly, computing the structural change is possible only after the RPM is constructed. To detect RPMs, we use *subgraph edit distance*, as we discuss below.

#### 3.2.1 Computing Reactant–Product Mapping

Intuitively, we should map the pair that is most similar to one another. We model this intuition using the idea of *subgraph edit distance*. Informally, the subgraph edit distance *sed(g, g′)* [14] is the minimum number of *edits* performed on *g* to convert it to some subgraph of *g′*. An edit is either addition or deletion of edges and vertices, or replacement of vertex or edge labels. We need to match *g* to all possible subgraphs of *g′* since in a decomposition reaction *AB ⟶ A + B, A* (and *B*) maps to a subgraph of *AB* and not the entire molecule. Delving into the subgraph space is necessary to accurately compute the structural change due to the reaction. We find the final set of RPM pairs using a greedy approach that chooses the reactant–product pair with the best *sed* first, the pair with the second best *sed* of the remaining pairs next, and so on until all products have been mapped. Formal definition of *sed*, as well as our algorithm to compute RPM, along with examples are presented in the Supplementary Methods.

### 3.3 Quantifying Structural Change

To quantify the changes due to the reaction, we first identify the *reaction centres*. Subsequently, we identify the *reaction signatures*, or the motifs we consider necessary for a reaction to occur.

#### 3.3.1 Reaction Centres

The reaction centre for an RPM pair (*A, B*) is the set of vertices in the product *B* to which new edges are added, or existing edges removed, during its transformation from *A*. For instance, consider the reaction in Fig 3a, and particularly focus on the pair (C00049, C00152). In this pair, the conversion involves a removal of the OH group and addition of the NH_2_ group. Thus, we have one reaction centre, which is the carbon atom attached to the bond involved with the change. The reaction centre is explicitly shown in Fig 3. The reaction centre for (C00020, C00002) pair is also shown in Fig 3. Note that we treat *OH* as a single vertex since *OH* chemically behaves as a single entity. Thus, the reaction centre is the vertex P and not O. Although it is more common to see one reaction centre in a pair, multiple reaction centres are possible.

**Figure 3:**
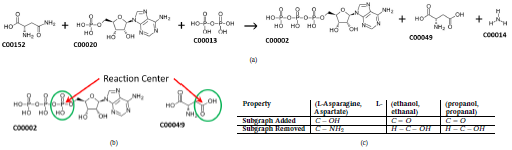
Illustration of reaction signature. (a) An example reaction, illustrating the conversion of the amino acid L-Asparagine (C00152) to L-Aspartate (C00049). The other reactants/co-factors in this reaction include ATP (C00002), AMP (C00020), Diphosphate (C00013) and Ammonia (C00014). (b) The reaction centres and signatures (green circle) in the reaction in (a). (c) The structural changes in the reactant–product pairs, in terms of subgraphs added and removed.

#### 3.3.2 Reaction Signature

The reaction centre only tells us the location of change. It does not necessarily tell us the reason, or the conditions necessary, for the change to occur. To predict pathways, we need to identify the conditions required for a reaction to happen. We build our prediction model based on the hypothesis that two molecules would undergo a similar change in a reaction if they contain a common “key” sub-structure that drives the forming or breaking of chemical bonds. Our hypothesis is motivated by the fact that many enzymes, such as alcohol dehydrogenases that convert alcohols to aldehydes, show a specificity towards the type of subgraph, *i. e*. sub-structure present in the reactants [16, 28]. Since the reaction centre is the location of the change, a straightforward approach would be to assign the reaction centre as this “key” subgraph. However, a single atom (or vertex) does not capture all of the atom-level interactions that take place. For instance, consider the reaction centre in L-Asparagine (C00152; see Fig 3a), which is a Carbon atom. Here, the Carbon is not only interacting with the *NH*_2_ group that gets replaced with the *OH* group, but also with the adjacent Oxygen and Carbon atoms. The strength of the *C = O* and *C* − *C* bonds, their charges, geometries etc. all play a role in the breaking of the *C* − *NH_2_* bond and its eventual replacement with *C* − *OH*. To generalise, the direct neighbours of the reaction centre influence the reaction. Based on this intuition, we define a *reaction signature, S(V_S_, E_S_)* as the immediate (‘one-hop’) neighbourhood of the reaction centre in the product of the (*A, B*) pair.

The reaction signatures of the two reactant–product pairs in the reaction in Fig 3a are shown in Fig 3b. It is easy to see that the reaction signature is a subgraph of the product. Note that when there are multiple reaction centres, there are multiple reaction signatures as well, where each signature represents the neighbourhood around the corresponding reaction centre. In general, the reaction centres identify the locations of change, and the reaction signatures encode the potential driving factor behind the change. Next, we formalise our mechanism to store the change itself.

#### 3.3.3 Detecting the change in (A, B) pair

Conceptually, in a pair (*A, B*), we want to store Δ = *B−A*, where Δ is the difference between the structures. Furthermore, given only *B* and Δ, we should be able to re-construct *A*. As we will see later, the ability to reconstruct the reactant *A* from just Δ and the product *B* lies at the core our of our algorithm’s ability to predict on unseen molecules. The reaction signature can change through either the addition or removal of subgraphs, as detailed in Supplementary Methods. To illustrate, Fig 3c shows the structural changes for three different pairs. The first pair is from Fig 3a. The other two pairs correspond to the reactions in Fig 2a. Notice that since both reactions in Fig 2a involve conversion of alcohol to aldehyde, their structural changes (along with the reaction centres and signatures, which are not shown in Fig 3c) are identical. The above illustration not only showcases how we capture structural changes in a reaction, but also demonstrates our precise ability to detect a common pattern among reactions. Armed with this technique, we next formulate the idea of a *reaction rule*.

### 3.4 Reaction Rules

Given a database of reactions 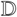, for each reaction 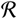, we identify all of its reactant–product pairs. From each pair (*A, B*), we extract and store the following information: (i) the reaction signature, (ii) the reaction centres, (iii) the subgraphs added and removed and (iv) all reactants in 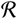 except *A*. These reactants are the co-factors or *helper* reactants that facilitate the reaction. For example, the oxidising agent would be stored as the helper reactant for (ethanol, ethanal) and (propanol, propanal) pairs in Fig 2a. For the (C00152, C00049) pair in the reaction in Fig 3a, both C00020 and C00013 would be stored.

We denote the above information, which is extracted from each (*A, B)* pair, as *L(A, B*), the reaction rule. Note that we do not store the pair (*A, B*) itself; we only store the structural change and its associated information. *L*(*A, B*) = *L*(*C, D*) if all of the four items listed above are identical. For example, *L*(*ethanol, ethanal*) = *L*(*propanol, propanal*).

To identify rules, we consider a support threshold *θ*, which decides the number of times a pattern of structural change must be seen, to be considered a reaction rule. Since novel pathway identification between rare molecules is of critical importance, we err on the side of exploration, and set the default *θ* = 1, which means any structural change is a pattern, even if it does not repeat across multiple reactions. The top ranking pathways can be manually screened at a later stage.

### 3.5 Pathway Prediction

We now discuss how the reaction rules described above can be employed to predict synthesis of a target product. A reaction rule serves two purposes: firstly, given any target product molecule, detect whether the rule is applicable on the molecule. If the rule is applicable, we must predict the reactants required to synthesize the given product. We introduce two graph operators: graph addition and subtraction, which enable the above. An example of the operations is shown in Fig 4a.

**Figure 4:**
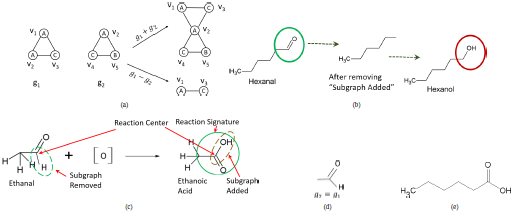
Illustration of graph operators and reaction rules. (a) An example of graph addition and subtraction. *υ*_*i*_ denotes the vertex IDs. (b) The process of predicting hexanol as the product pair of hexanal. (c) Features of the rules extracted from the conversion of ethanal to ethanoic acid. (d) *g*_2_ and *g*_1_ corresponding to the example RRN discussed in the main text. (e) Structure of hexanoic acid.

**Graph Addition.** *m*(*V_m_, E_m_*) = *g*(*V, E*) + *g′*(*V′, E′*). The resultant graph *m* has *V_m_* = *V* ∪ *V′, E_m_* = *E* ∪ *E′*. For *m* to be connected, it must satisfy that *V* ∩ *V′* ≠ 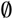.

**Graph Subtraction.** *m*(*V_m_, E_m_*) = *g*(*V, E*) − *g′*(*V′, E′*). The resultant graph *m* has *V_m_* = {*υ* ∈ *V, υ* ∉ *V′*}, *E_m_* = {*e* = (*υ*_1_, *υ*_2_) ∈ *E*| *υ*_1_, *υ*_2_ ∈ *V_m_*}.

Algorithm 1 presents the pseudocode of applying a reaction rule. Let *B* be a target product and *L* be the reaction rule that we want to apply on *B* if chemically feasible, i.e. if the rule is applicable, based on the presence of appropriate subgraphs. Recall our hypothesis that the presence of the reaction signature is the cause of the reaction. Second, due to the reaction, the “Subgraph Added” of *L* gets attached at the reaction centre *c*. Thus, we first merge the reaction signature with the “Subgraph Added” to create a single merged graph *m*. If *m* is a subgraph of *B*, then *L* is applicable on *B*. If the check passes, we proceed to the next step of formulating the reactants that can synthesize *B*. Since the reaction centre is present both in the signature and the “Subgraph Added”, *m* is guaranteed to be connected.

#### Algorithm 1: *ApplyRule* (*L, B*)

~~~
1: *m* ← *L.signature* + *L.subgr aphAdded*
2: **if** *m* с *B* **then**
3:    *B* ← *B* − *L.subgr aphAdded*
4:    *A* ← *B* + *L.subgr aphRemoved*
5:    **return** {*A, L.helper Reactants*}
6: **else**
7:    **return** Not applicable
~~~

First, we construct the reactant pair of *B* using *L*. We remove the “Subgraph Added” from *B* (line 3) and then merge the “Subgraph Removed” component with *B* to create the reactant pair *A* (line 4). Finally, the helper reactants in *L* are fetched and their reaction with *A* is predicted to synthesize *B* (line 5). Note that neither *B* nor *A* is required to be present in the training database — only a matching subgraph need be present.

To illustrate, let us revisit our original example of synthesising hexanal from the training database in Fig 2a. The structure of hexanal is shown in Fig 4b. *L*(*ethanol, ethanal*) is a reaction rule, which also occurs again in *L*(*propanol,propanal*). The reaction signature for this rule is the subgraph within the green box in Fig 2aa and the reaction centre is the central Carbon atom participating in the C=O bond. The “Subgraph Added” and “Subgraph Removed” correspond to columns 2 (and 3) of Fig 3c and the helper reactant contained in this rule is the oxidising agent [O]. The reaction rule is applicable on hexanal since the merged graph of the signature and “Subgraph Added” is a subgraph of hexanal. Now, to identify the reactant pair, first we remove “Subgraph Added” and then attach the OH at the reaction centre. Consequently, we generate the hexanol molecule. The step-by-step process is shown in Fig 4b. Finally, we predict hexanol +[O] as the reaction since [O] is the helper reactant stored in the rule. Thus, we are able to predict the synthesis of hexanal from hexanol even though we have not seen either of the molecules in the training database.

### 3.6 Reaction Rule Network (RRN)

While we have described above, the procedure to predict a reaction that could synthesize a target molecule (also see Algorithm 1), our goal is to predict pathways — essentially a chain of reactions. Furthermore, between a source and a target molecule, there could be hundreds of pathways. How do we identify and rank only the top-*k* best paths? To overcome these challenges, we propose the idea of a *reaction rule network (RRN)*.

Each node in the RRN corresponds to a reaction rule, and we want to ensure the following property: if there exists a pathway *P* = {*R*_1_, ⋯, *R_n_*} from molecule *A* to *B*, such that reaction *R_i_*; happens through rule *L_i_*, then, there should be a path from *L_n_* to *L*_1_ in the RRN. Towards that goal, we notice that rules *L*_1_ and *L*_2_ can be applied consecutively if the product of *L*_1_ is a reactant in *L*_2_. In such a case, we should have a directed edge from *L*_2_ to *L*_1_. However, neither the product nor the reactant may be present in the database. We need to capture this dependency between *L*_1_ and *L*_2_ only from the structural change information that we store. To capture all of these properties, we formally define the *RRN* as follows:

*Definition 3.1*. REACTION RULE NETWORK (RRN). Let 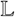 be the set of all rules mined from our training database. The RRN *N*(*V_N_, E_N_*) is a directed graph where *V_N_* = 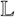. Let *g*_2_ = *L*_2_.*signature* − *L_2_.subgraphAdded*+ *L_2_.subgraphRemoved* and *g*_1_ = *L*_1_.*signature*+ *L*_1_.*subgraphAdded* and *e* = (*L*_2_, *L*_1_) ∈ *E_N_* if *g*_1_ ⊆ *g*_2_.

In the above definition, *g*_2_ is the subgraph that must be present on any reactant on which *L*_2_ is applicable. On the other hand, *g*_1_ is the subgraph that must be present on any product generated through *L*_1_ (follows from Algorithm 1). Thus, if *g*_2_ is a (subgraph isomorphic) subgraph of *g*_1_, then the product of *L*_1_ can feed in as a reactant to *L*_2_. To illustrate the RRN, consider a training database where in addition to the two reactions in Fig 2aa, we also have the oxidation of ethanal to ethanoic acid shown in Fig 4c. Furthermore, we consider every unique structural change as a pattern. Thus, there are two reaction rules; rule *L*_1_ corresponding to the conversion of alcohol to aldehyde, and rule *L*_2_ corresponding to the conversion of ethanal to ethanoic acid. The reaction signature, subgraph added and subgraph removed for *L*_2_ is also shown in Fig 4c. The resultant *g*_2_ and *g*_1_, as shown in Fig 4d, are isomorphic, and consequently, there is an edge from *L*_2_ to *L*_1_ in the resultant RRN.

The formalization of the RRN completes the offline model building component. Next, we discuss the online query (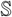, *T*), where the goal is to find a pathway from *A* to *T* where *A* ∈ 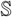 is one of the source molecules.

#### Algorithm 2: *TopKPaths*(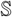, *T, k*)

~~~
**Input:** Set of source molecules 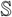, Target molecule *T* and number of paths *k*
**Output:** Top-*k* paths ranked according to heuristic *H_d_*
1: *Initialise PQ* ← *ϕ*
2: *InitialiseanswerSet* ← **new** *MaxHeap*(*k*) /* Max heap of size k ordered by dist */
3: *srcVertices* ← {*υ* ∈ *V_N_*\*υ*.*signature* + *υ.subgraphAdded* ⊆ *T*}
4: **for** υ ∈ *srcVertices* **do**
5:   *L* ← **new** *PQNode*()
6:   *L.path* ← {*υ*}
7:   *L.reactant* ← **ApplyRule**(*υ, T*)
8:   *L.dist* ← *H_d_*(*υ, L.reactant*, 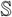)
9:   *L.pathway* ← {*L.reactant*}
10:  *PQ* ← *PQ* ∪ *L*
11: **while** *PQ* ≠ *ϕ* **do**
12:  *L* ← *PQ.extractMinDist*()
13:  **if** *L.reactant* ∈ 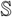 **and** (*|answerSet|* < *k* **or**
     *L.dist* < *answerSet.Top*().*dist*) **then**
14:    *answerSet* ← *answerSet* ∪ *L.path*;
15:  **for** *L_adj_* ∈ *N.Adj*(*L.path.lastVertex*) **do**
16:    *M* ← **new** *PQNode*()
17:    *M.reactant* ← **ApplyRule**(*L_adj_, L.reactant*)
18:    *M.path* ← *L.path* ∪ {*L_adj_*}
19:    *M.pathway* ← *L.pathway* ∪ {*M.reactant*}
20:    *M.dist* ← *Hd*(*L_adj_, M.pathway*, 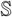)
21:    **if** *|answerSet|* < *k* **or** *M.dist < answerSet.Top*().*dist*
       **then**
22:      *PQ* ← *PQ* ∪ *M*
23: **return** *answerSet*
~~~

### 3.7 Answering Queries on the RRN

To illustrate our query answering strategy, we continue with the RRN outlined above. Suppose the query is to find a pathway from hexanol to hexanoic acid (Fig 4e). Note that neither of the query molecules are in the reaction database. We initiate by searching for a rule that is applicable on the target molecule, hexanoic acid. In our two-node network, rule *L*_2_ is applicable (line 1 in Algorithm 1). On applying *L*_2_ on hexanoic acid, hexanal is generated as the reactant pair. Since hexanal is not the source molecule, we continue searching by applying the adjacent rule *L*_1_. Since *L*_1_ is connected from *L*_2_, we are guaranteed that *L*_1_ is applicable on the reactant produced by *L*_2_, which is hexanal. On applying *L*_1_ on hexanal, hex-anol is generated as the reactant pair, which completes the query since it is the source molecule. The resultant pathway is therefore hexanol 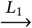 hexanal 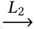 hexanoic acid.

To generalize the above strategy, we first identify nodes (or rules) that are applicable on the target molecule. From each of these rules, a reactant is generated. If the reactant is one of the source molecules then we stop. Otherwise, we continue exploring each possible path using breadth-first search (BFS) either till all paths are exhausted or a source molecule is reached. We call this strategy the *BFS exploration*. Exploration using *BFS* guarantees that the first pathway found is the shortest, in terms of length. The exploration algorithm can easily be generalized to find the *k* shortest paths as well. While *BFS* is simple, it is often not scalable in a large RRN due to the large number of paths that exists. Furthermore, the BFS strategy does not use the knowledge of the source molecule to optimise the searching process. To overcome these weaknesses, we explore an alternative algorithm, based on *best-first search* [32].

#### 3.7.1 Heuristic H_d_: Minimizing structural changes in every step

We hypothesize that nature avoids reactions that cause drastic alterations to the structure of the reactant. This can also be appreciated in terms of the enzymes — enzymes are highly specialized and perform an incremental structural change to a substrate, rather than wholesale structural changes. We model this effect through a distance function that minimizes the total structural change in a pathway, in addition to minimizing the distance to a source molecule *A*. Specifically, the optimization function at a specific pathway *P* = {*X*_1_,…,*X_n_* } of *n* molecules (*n* − 1 reactions) minimizes the function below:

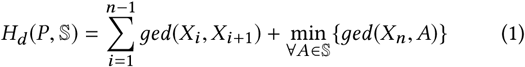
where *ged(g, g′)* is the *edit distance* between graphs *g* and *g′* [34]. Edit distance between two graphs is defined analogously to subgraph edit distance. Specifically, it is the minimum number of edits required to convert *g* to *g′*. The primary difference with *sed(g, g′)* is that *g* is converted to *g′* instead of a subgraph of *g′*. Consequently, *ged(g, g′)* is symmetric. Based on *H_d_*(*P*, 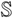), we optimise search paths using best-first search, as listed in Algorithm 2.

Algorithm 2 explains how we we optimise search paths using best-first search, based on *H_d_*(*P*, 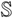). We initialise a priority-queue (PQ) that orders rules in ascending order of their distance and the answer set (AS) as a max heap of size *k* (lines 1–2). First, we insert all applicable rules on the target molecule *T* in the PQ (lines 4–10). Next, we pop the top rule *L*, i. e. the rule that generates a pathway minimising Eq 1 and check if any of the source molecules has been reached. If it has, then we insert the path in the answer set only if the distance (of *L*) is lower than the distance of the *k^th^* best path identified so far (lines 13–14). Otherwise, we extract each of *L′s* adjacent rules, generate the resultant reactants, and insert the rules based on their distance in the PQ (lines 15–22). In this step, we insert a rule in the PQ only if the distance is lesser than that of the *k^th^* best path (lines 21–22). We again pop the next applicable rule with the lowest distance and this process continues until all paths to the source molecules are exhausted (line 11).

The details of the 20 pathways can be found in Supplementary Table S2.

### 3.8 Datasets used

We used the KEGG [15] as our training database. KEGG harbours an extensive collection of >10,000 biochemical reactions known to occur in various organisms. We also obtained organism-wise reaction sets, where each set is a subset of the KEGG, containing the known reactions in a single organism, such as yeast, *E. coli*, etc. In total, we have 2,641 organisms, which results in 2,641 reaction sets. These datasets were obtained via Path2Models [1], which is based on KEGG. Our algorithms are implemented in Java JDK 1.7.0 and evaluated on a PC with 12GB memory and Intel i5 2.60GHz quad core processor running Ubuntu 13.04.

## 4 RESULTS

In this section, we establish that our pathway predictions are accurate, and that the proposed technique is scalable to large reaction databases. Ours is the first technique that is fully automated, can answer queries on unseen molecules, and requires no information other than the structure of the molecules. Due to this simplicity of our technique, we are the first to scale to a database as large as 150,000 reactions.

Our major results are four-fold. First, we query on those source and target molecules present in the training database. The presence of query molecules in the training set is enforced only to allow us to compare the performance with the state-of-the-art pathway prediction techniques such as *RouteSearch* [20] and *MRE* [19]. We demonstrate how our heuristic *H_d_* picks up natural biosynthetic pathways very frequently, much more than other state-of-the-art methods. Second, we show that the common biosynthetic pathways are optimised across organisms in nature. Third, we remove the constraint of requiring the source molecules in the training database and show that we predict viable retrosynthetic pathways for known and new molecules. Finally, we show that our results are accurate, by means of cross-validation, and that our algorithm can scale well for very large reaction databases.

### 4.1 *H_d_* consistently picks up natural pathways with high probability

In any pathway prediction algorithm, all predicted pathways are ranked according to some score, and finally the top-k highest scoring paths are studied further for feasibility. Ranking the predicted pathways is very important since there are often hundreds of paths between two molecules, and a high rank should signify high biochemical plausibility. As discussed earlier, we use Eq 1 as the ranking function in our algorithm. To benchmark, we choose 20 pathways involved in the biosynthesis of amino acids and important precursors in central carbon metabolism, similar to those used in [4].

For the selected pathways, we predict by querying using their source and target molecules and extract the top 10 predicted paths. The training database for this experiment corresponds to the reaction set of *E. coli*. Table 1 presents the rank of the actual pathway by each of the techniques. As clearly evident, the actual pathway consistently ranks among the top 10 in our algorithm, while being mostly absent in RouteSearch. MRE is able to predict only 10 of the 20 pathways. Although MRE occasionally ranks the correct result higher than our method, it clearly lags behind our method in the overall head-to-head comparison (4-14 with 2 ties). These results point towards the superior ability of our technique to identify pathways reliably, and also rank the biologically favoured pathways much higher.

**Table 1:**
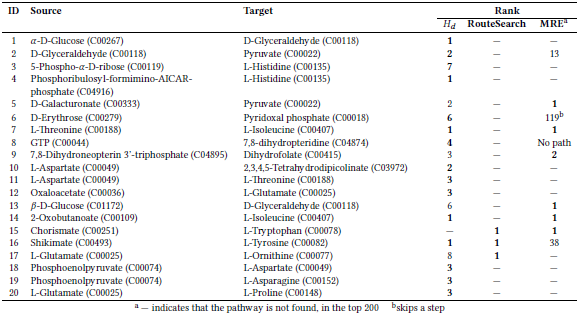
Pathway Prediction comparison of our algorithm (specifically, using the heuristic *H*_d_) versus RouteSearch and MRE. The source and target molecules are indicated along with their KEGG CIDs. Bold-faced rank displays the winning algorithm for each row.

### 4.2 Nature appears to optimize pathways across organisms

Although the performance of our technique is clearly superior, the native biosynthetic pathway did not always rank the highest. We now investigate the possible reason behind this behaviour. We base our investigation on the hypothesis that nature prefers common pathways that are feasible across multiple organisms instead of a single organism [10, 27]. To test this hypothesis, we again considered the 20 commonly occurring pathways from central carbon metabolism and amino acid synthesis, which are common to many organisms. We performed the pathway prediction on the organism-specific reaction sets and computed the rank of the actual pathway using our heuristic *H_d_*. We then computed an *Aggregate Score* for each path, as 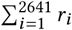, where *r_i_* is the rank of the actual path in organism *i*. The pathways are then globally ranked based on their Aggregate Score; the lower the Aggregate Score, better is the rank (see Table 2). We also list the total number of unique pathways that exist between the source and the target molecules to fully expose the complexity of the prediction and ranking task. The results clearly reveal that nature can be better explained when the analysis covers multiple organisms than a single one. More specifically, the actual pathway consistently ranks highest in 60% of the paths and within the top three for more than 90% of the paths. Nature’s preference to use the same pathway for biosynthesis of these molecules across all organisms is also captured by the *Average Rank* column in Table 2.

**Table 2:**
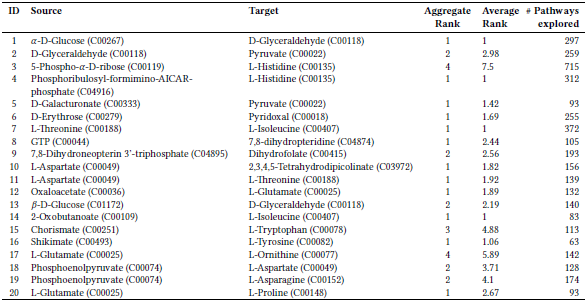
Pathway prediction results across multiple organisms. The source and target molecules are indicated, along with their KEGG IDs. Also shown are the aggregate rank and average rank (across all organisms), along with the total number of pathways explored. The details of the 20 pathways are given in Supplementary Table S2.

### 4.3 Retrosynthetic predictions compare favourably with other methods

In addition to the pathways we outlined above, we here show that we perform comparably or better than MRE, in nearly all retrosynthesis examples discussed in [19]. We predict retrosynthesis pathways for commercially important metabolites, such as itaconate, naringenin, 1,3-propanediol, xylitol etc. We find that in a majority of cases, we are able to recover known pathways or predict shorter biologically plausible pathways for retrosynthesis. We summarise our retrosynthesis predictions in Table 3, alongside comparisons with MRE/FMM.

**Table 3:** Retrosynthetic pathways to various molecules, as predicted by our algorithm, in comparison with other methods such as MRE and FMM.

| # | Source | Path (intermediates)<sup>a</sup> | Target | Comments |
| --- | --- | --- | --- | --- |
| 1 | Pyruvate (C00022) | C01011 → C0531 | Itaconate (C00490) | Same as FMM |
| 2 | L-Tyrosine (C00082) | C00811 → C00223 → C06561 | Naringenin (C00509) | Same as MRE/FMM |
| 3 | L-Tyrosine (C00082) | C00811 → C00223 | Resveratrol (C03582) | Same as MRE/FMM |
| 4 | Glycerol (C00116) | C00969 | Propane-1,3-diol (C02457) | Same as MRE/FMM |
| 5 | Glycerol (C00116) | C00577 → C00546 → C00937 | (R)-Propane-1,2-diol (C02912) | Also predicts two paths shorter than MRE, in addition |
| 6 | D-Xylose (C00181) | — (direct) | Xylitol (C00379) | Shorter than MRE; same as FMM |
| 7 | HMG-CoA (C00356) | C00418 → C01107 → C01143 →<br>C00129 → C00448 → C16028 | Artemisinic acid (C20309) | Identical pathway as in MRE, but differs from [31]; paths from acetyl-CoA are more convoluted |
| 8 | Chorismate (C00251) |  | L-Tryptophan (C00078) | Unable to find relevant paths |
| 9 | 5-Phospho-α-D-ribose (C00119) | C02739 → C02741 → C04896 →<br>C04916 → C04666 → C01267 →<br>C01100 → C00860 | L-Histidine (C00135) | Recovers native path in top 10, amongst other probable paths; MRE does not recover native path but proposes shorter routes |
| 10 | Pentachlorophenol (C02575)<sup>b</sup> | C03434 → C07099 → C07097 →<br>C06329 | 2-Maleylacetate (C02222) | Identical pathway to [2] |
| 11 | Atrazine (C06551)<sup>b</sup> | C06552 → C06553 → C06554 →<br>C06555 | Urea-1-carboxylate (C01010) | Same as MRE |

<sup>a</sup>Intermediates are indicated only by KEGG CIDs, for brevity
<sup>b</sup>these are xenobiotic degradation pathways, rather than retrosynthetic pathways

For itaconate, an important value-added precursor from biomass [33] we recovered the same path as predicted by FMM. For production of naringenin, an important plant secondary metabolite, and resveratrol, we find the same pathway identified by MRE and FMM. For the production of xylitol, our top-ranked pathway is shorter than that proposed by MRE, and agrees with FMM. For artemisinic acid, an important anti-malarial drug, synthesised in metabolically engineered *S. cerevisiae* [31], we were able to predict the same path as MRE, from HMG-CoA, although this differs from [31]. For paths from acetyl-CoA to artemisinic acid, and chorismate to L-Tryptophan, the top ranked paths from our algorithm are not very relevant, perhaps due to the occurrence of very high-degree metabolites, such as acetyl-CoA and pyruvate.

Furthermore, we also examined some of the pathways evolved by organisms to degrade anthropogenic chemicals such as pentachlorophenol [2, 8]. We find that we are able to generate the identical pathway between pentachlorophenol (C02575) and Maleylacetate (C02222), as indicated in Table 3. It is interesting to note that this predicted pathway is one of several possible pathways, given that we can apply many reaction rules to every intermediate. We also find that MRE and FMM are unable to find any pathways between these compounds, illustrating the importance of our ability to generalise reaction rules, as well as handle novel molecules. MRE and our approach both correctly predict another pathway where atrazine (C06551) is converted to urea-1-carboxylate (C01010). Together, these results illustrate the ability of our approach to not only predict retrosynthetic pathways, but also possible pathways that organisms may use to metabolise xenobiotics. Importantly, our heuristic of minimising the metabolic transformations in a reaction enables us to recover the very pathway these organisms have evolved to breakdown xenobiotics.

### 4.4 Cross-validation illustrates the high accuracy of our pathway predictions

First, we evaluate through 5-fold cross-validation. Specifically, we split the KEGG Dataset into five parts, learn the training model on four parts and predict on the fifth part. This process is repeated to cover each part as the test set. For our prediction query, we pick arbitrary pathways from the test set and check if the exact pathways are predicted. We always ensure that the source and the target molecules are not part of the training set. Fig 5a presents the prediction accuracy against the training dataset size. To understand the results better, we segregate them into pathways of length 1, 2, and ≥ 3. The trends are similar across all lengths and the results saturate at around ≈35,000 reactions in the training dataset. As expected, the accuracy is better for single length pathways since the search space is smaller. Theoretically, the search space increases exponentially by a factor of *d* with each hop, where *d* is the average degree of the RRN. For all three pathway lengths, the accuracy is higher than 80% at ≈35,000 training reactions and beyond.

**Figure 5:**
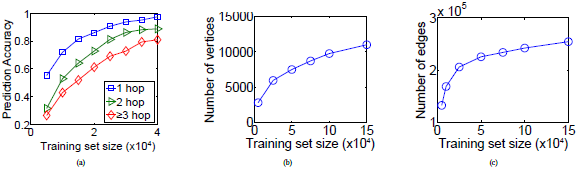
(a) Accuracy of pathway prediction against training dataset size, for pathways of varying lengths. (b-c) Effect of increasing training set size on (b) the number of vertices and (c) the number of edges in the RRN.

We continue with a similar line of analysis and next check how the size of the RRN saturates with training dataset size. Figs. 5b and 5c present the results. The number of edges saturates quicker than the number of vertices. This means that although new rules are discovered on increasing the training size, these rules are outliers and cannot be used in sequence with other rules. Notice that just like accuracy, the number of edges saturates at around 40,000 reactions as well. This correlation is not surprising.

### 4.5 Scalability

In this section, we benchmark the scalability of our technique. As we have pointed out, this is the first technique to move beyond individual case-studies and scale to thousands of reactions. First, we investigate the querying time in our 5-fold cross validation study against the training dataset size. In these experiments, we study the running time for both the heuristic search *H_d_* defined in Eq 1, as well as the basic breadth first search. To understand the results better, we plot the running time for pathways of length 1, 2 and ≥ 3 separately. Figures 6a–c present the results. An interesting pattern emerges from these results. For short pathways, BFS is faster than the heuristic of minimizing structural changes. This is expected since BFS blindly applies all rules within 1 or 2 hops. However, as the length of the pathway grows, the number of possible paths grows exponentially. Hence, BFS finds it hard to connect to the source amid so many possibilities. This pattern is even more evident in Fig 6d, where we study the growth rate of querying time with pathway length. While BFS is competitive till length 2, beyond that, it is not scalable.

**Figure 6:**
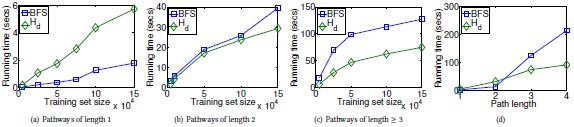
(a-c) Variation in querying time against the size of the training set, for pathways of length 1, 2 and ≥3, (d) Variation in querying time against the length of pathway. Both BFS (blue squares) and the best-first search using the *H_d_* heuristic (green diamonds) are depicted.

While the length of the pathway is one factor, another dominant factor in querying time is the number of signatures present in the target product. If the target contains large number of signatures, then more number of rules are applicable on it. Consequently, the search space increases and the querying time grows. This effect is clearly visible in Fig 7a.

**Figure 7:**
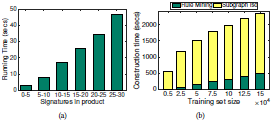
(a) Variation in querying time against the number of reaction signatures present in target molecule. (b) Effect of increasing training set size on the time to build our prediction model.

Fig 7b shows the growth rate of construction time against the training dataset size. We partition the time taken to build the network into two components. The first component looks at the time taken to mine the rules and the second component checks the time taken to build the network from these rules. As visible, the majority of the model building time is spent on constructing the network. To build the network, we need to compare all pairs of rules. Each of these comparisons involves subgraph isomorphism tests and hence it takes more time than rule mining, which is a linear scan across all reactions in the dataset. The overall growth of the construction time decreases beyond 75,000 reactions since at this point the number of rules mined (Fig 5b) also saturates.

## 5 DISCUSSION

Is it possible to synthesise molecule B from molecule A? Are there alternative pathways to synthesise a molecule, other than the one followed by cells of living organisms? Why do organisms in nature choose a particular pathway to synthesise a metabolite, say pyruvate, from glucose? In this paper, we have developed a pathway prediction technique that can answer these questions. The proposed system is the first fully-automated technique that can operate at the level of hundreds of thousands of reactions and answer queries in seconds. This level of sophistication is achieved through a graph mining based approach, which automatically mines cause-and-effect patterns of structural transformations from a training database of chemical reactions. These patterns are employed to construct an abstract representation of the reaction space in the form of a RRN. This abstract representation lies at the core of our ability to make rapid predictions, even on molecules that we have never seen before.

Many earlier studies have approached path finding in metabolic/chemical reaction networks; however, they typically fall short in one or more of the following: (a) they rely on the existence of query molecules in their database, or (b) their pipeline involves the application of hand-curated rules such as atom–atom mapping information, or (c) they only work for specific classes of reactions (also see. Using no more information than the molecular structure of every molecule in the reaction database, we have developed a powerful pipeline for predicting pathways between any two metabolites.

Our key findings fall into three categories. First, we have an efficient reactant–product mapping that is built on subgraph edit distance. It enables us to accurately track changes in chemical moieties across the entire spectrum of biochemical reactions. Next, we identified reaction signatures, which are essentially subgraphs necessary for the reactions to occur. Next, we embedded information about the *reaction centres* in a given metabolic network onto another network, the RRN. This novel representation enables us to predict a series of reactions (or, a pathway) connecting two metabolites, which may not even belong in the original reaction database.

We then proceeded to ask a more fundamental question about the organisation of metabolic networks: What is the key underlying design principle of known metabolic pathways? For example, it is well-known that standard biochemical pathways do not represent shortest paths in the network — there are likely other constraints such as energetics in play. Other studies [27] have shown that central carbon metabolism is a minimal walk between key precursor metabolites. We have here shown that across an assortment of pathways, nature appears to minimize the incremental biochemical change occurring, from the reactant to product, in every step of the reaction. By employing a heuristic built on this logic, we correctly recover a majority of pathways (see Table 2) from carbohydrate, amino acid and fatty acid metabolism. In certain cases, we observe that a different pathway is in use by nature, clearly owing to energy considerations. For example, the path from D-Mannose to L-Galactose in nature may be convoluted, owing to energy considerations: D-Mannose → GDP-mannose → GDP-L-galactose → *β*-L-Galactose → L-Galactose, even though a simple epimerisation reaction may theoretically be possible. It is important to note that our graph formalism, coupled with our heuristic has enabled us make reliable predictions, even in the absence of important information such as atom–atom mapping or A*G* values for different reactions.

We have also predicted retrosynthetic pathways to commercially important molecules such as 1,3-propanediol, naringenin, itaconate and artemisinic acid, and we compare favourably with previous methods such as MRE [19] and FMM [7]. Importantly, we are able to additionally predict pathways for compounds such as pentachlorophenol, which MRE and FMM are unable to. Our method also enables us to predict pathways for compounds not present in the training database.

Finally, we also demonstrated that our approach is very scalable. This is particularly important in the light of the fact that many studies have pointed out that our current understanding of microbial metabolism is rather myopic — many more organisms from diverse phyla need to be reconstructed, and even for many current metabolic network reconstructions, major gaps in the reactome are present [25]. A comparison with the BRENDA enzyme database also showed that only a third of the enzymatic activities in BRENDA are covered by currently available metabolic networks [25]. Given the significant imminent expansion in metabolic network databases, a scalable approach such as ours bears special significance. By synthetically expanding the KEGG database to about 150,000 reactions, we show that our approach is still very fast, able to answer queries in a matter of seconds.

Our method is not without limitations. In choosing to keep the input information as minimal as possible, to enable widespread applicability, we have chosen to leave out thermodynamics from the picture, often very essential for accurate predictions and ranking of pathways. Nevertheless, we demonstrate that even without thermodynamic information, we are able to recover a majority of natural biosynthetic pathways. Further, it is often difficult to obtain accurate measurements of changes in free energy, especially those which are organism-specific. Also, like most other similar approaches to predict reactions, the accuracy of our approach is limited by the accuracy of the reaction database, KEGG, in this case. KEGG also contains no information about the reversibility of reactions, and essentially assumes all reactions are reversible. However, it will be possible to integrate information from other databases such as MetaCyc, or even use a consensus; the scalability of our algorithm will be particularly handy in such scenarios.

## 6 CONCLUSION

In sum, we see three major contributions of our study. First, we define a robust reaction–product mapping method using subgraph edit distance, which is fast and reliable. This enables us to construct a novel representation of a database of chemical reactions in terms of a RRN that lends itself to rapid querying for pathways to synthesize even molecules that are not present in the original reaction databases. Next, we define a heuristic to perform searches on this network, by minimizing the extent of transformation in every reaction. Searching using this heuristic very effectively recovers known native pathways across organisms, and enables a realistic ranking of predicted alternate biosynthetic pathways. Finally, we demonstrate the ease with which we can provide solutions to retrosynthesis queries. Importantly, our approach uses no information other than the chemical structure of the molecules in every individual reaction, and yet gives very accurate results and scales up to over a hundred thousand reactions.

## ACKNOWLEDGMENTS

The authors thank Pankaj Kumar and Aarthi Ravikrishnan for useful discussions and Manikandan Narayanan for a critical reading of the manuscript. This work was supported by the Indian Institute of Technology Madras grant CSE/14-15/5643/ NFSC/SAYN to SR.

## SUPPLEMENTARY METHODS

Refer to the supplementary file.

